# Pro-GAT: Reconnecting Fragmented PROTACs Using Graph Attention Transformer

**DOI:** 10.64898/2026.02.22.707266

**Authors:** Saahithi Vemuri, Laxmi Priya Bijigiri, Sanjana Gogte, Vani Kondaparthi

## Abstract

PROTACs work by bringing together a protein-of-interest ligand and an E3 ligase recruiter to trigger targeted degradation. However, Diffusion-based generative models frequently produce chemically invalid or disconnected linker structures that satisfy global geometric constraints but violate local bonding requirements. These models operate in continuous coordinate space and therefore lack explicit mechanisms for enforcing discrete chemical connectivity under fixed-anchor constraints. Invalid, disconnected outputs recur rather than being a rare exception, such that naive resampling is not an effective method to obtain valid chimeras.

Pro-GAT is a graph attention-based framework for geometry-preserving molecular graph repair, capable of functioning on chemically disconnected diffusion-generated PROTAC candidates by predicting bounded coordinate corrections and constrained atom-type modifications using geometry-aware graph attention network (GAT) layers. The proposed model is trained on PROTAC datasets with added disconnections to overcome systematic connectivity failures in diffusion-based PROTAC generation with fixed anchors. When combined with DiffPROTACs and DiffLinker, Pro-GAT improves the percentage of chemically valid candidates in the aggregated output from 76.70% to 83.92% and 63.16% to 68.73% while maintaining 80.18% and 63.80% uniqueness levels of valid candidates respectively, thus facilitating the generation of usable PROTAC candidates from invalid diffusion samples. Pro-GAT was used in a case study of the 7Z76 ternary complex to repair DiffPROTACs and DiffLinker generated samples, which gave rise to connected chimeras whose docking scores were comparable to the original 7Z76 structure.

## 1. Introduction

PROTACs are heterobifunctional molecules that induce targeted protein degradation by simultaneously binding a POI and an E3 ubiquitin ligase, forming a ternary complex that elicits ubiquitination and proteasomal degradation [1]. In contrast to traditional small-molecule inhibitors, PROTACs confer unique advantages toward ‘undruggable’ targets: enabling sub-stoichiometric degradation, overcoming resistance mutations, and making intracellular protein homeostasis accessible [2, 3]. More than 20 clinical trials have advanced for PROTACs in oncology, neurodegeneration, and immunology; molecules such as DT2216, the BCR-ABL degrader, have shown clinical efficacy [4-6].

Despite this promise, PROTAC linker design remains a central challenge in targeted protein degradation [7-9]. Linkers represent the primary degree of freedom governing PROTAC efficacy, as they must span precise POI-E3 distances, support stable ternary complex formation, and satisfy physicochemical and synthetic constraints [10, 11]. Diffusion-based generative models sample linker atom coordinates under fixed ligand anchors, but do so without enforcing discrete bonding constraints [12]. However, these samples frequently appear disconnected, failing local bonding constraints despite being close to feasible solutions. Naive oversampling does not resolve this issue, as disconnected structures recur systematically rather than as rare sampling noise [12]. Increasing the number of generations not only incurs substantial computational cost due to the large number of diffusion denoising steps per sample, but also shifts sampling toward low-probability regions of the distribution, increasing the fraction of structurally implausible samples, reducing uniqueness among valid generations, and discarding many geometrically promising but chemically disconnected candidates [13-15]. Consequently, this creates a gap between efficient geometric exploration and chemical realizability that cannot be addressed by generation alone.

We introduce Pro-GAT, a graph attention–based repair model operating on the diffusion outputs which are chemically invalid due to disconnected fragments of the linker. Pro-GAT predicts bounded atomic coordinate corrections together with atom type retention or modification decisions to restore connectivity across broken linker segments while maintaining the three-dimensional geometry generated by diffusion. Trained on augmented PROTAC datasets [16] with synthetic disconnections, Pro-GAT learns context-aware local repair operations enabling realistic bond formation without performing global conformational optimization.

## 1. Methodology

### 1.1. Pro-GAT Overview

Pro-GAT operates on PROTAC linker candidates generated by diffusion-based models. It operates on diffusion outputs with disconnected linker fragments and learns to restore covalent connectivity by predicting bounded coordinate adjustments and constrained atom-type changes.

Pro-GAT enhances the usable yield of the models by recovering a significant fraction of diffusion outputs that are chemically invalid due to linker disconnection, without retraining or modifying the underlying generator.

Pro-GAT is intentionally limited to a local, geometry-preserving repair model. It predicts bounded coordinate adjustments and atom-type changes in order to recover covalent connectivity in disconnected linker structures without ever inserting new atoms or extending linker length. Following repair by Pro-GAT, all successfully repaired molecules are sanitized according to RDKit conventions, which enforce the typical chemical valence rules and correct residual valency violations at reconnection. The outcome is that the repaired output of the pipeline is guaranteed to be chemically valid and RDKit-sanitizable, whereas molecules that cannot be resolved by connectivity repair and subsequent sanitization are excluded.

Pro-GAT cannot repair structures where the spatial separation between ligand attachment points exceeds that which can be spanned by the linker backbone [7-9, 17]. In this case, no reassignment of atom types and/or bond orders, whether during the repair or subsequent sanitization, can yield a chemically connected structure and regeneration with a longer linker is required rather than post hoc repair.

### 1.2. Dataset Construction

The dataset for Pro-GAT is prepared based on the existing benchmark of DiffPROTACs [12] and it’s published supplementary data. DiffPROTACs provides ligand pairs, linker-length annotations, and pre-defined train-validation-test splits that are all kept unchanged so that each ligand pair in the repair data belongs to the same split as in the original benchmark.

For each PROTAC in the DiffPROTACs training set, we carry out diffusion inference to generate 10 samples per ligand pair under the same conditions as in the original evaluation protocol. Training supervision is restricted to PROTACs with short linker lengths (5–8 atoms), for which reliable connected reference structures are available in the DiffPROTACs supplementary data. It is empirically found that including supervision targets from longer linker lengths during training degrades repair performance on short linkers. This likely follows from the increased structural and geometric heterogeneity associated with long flexible linkers. In contrast, training exclusively on short linker lengths yields stable repair behaviour in the short-linker regime while also generalizing well to longer linker lengths at inference time.

When this diffusion inference gives disconnected or chemically invalid linker structures in a 5-8 linker-length regime, we retrieve the corresponding connected PROTAC structures from the supplementary dataset, select the reference structure whose linker geometry is most similar to the disconnected diffusion output, and use them as supervision targets. This procedure yields paired training examples of the form ‘disconnected diffusion output → connected valid structure’, enabling Pro-GAT to learn locality-preserving corrections for common diffusion failure modes. Since all supervision targets are drawn from DiffPROTACs’ own valid outputs, Pro-GAT operates strictly within the chemical space already explored by the generator, focusing on enforcing chemical realizability rather than expanding generative diversity.

### 1.3. Model Architecture

Pro-GAT takes as input a disconnected PROTAC graph with 3D coordinates, passes it through stacked attention-based message-passing [18] layers to produce refined node embeddings, and then uses three different MLP heads [19, 20] (https://scikit-learn.org/stable/modules/neural_networks_supervised.html) at each node to predict the corrected atom type, a coordinate adjustment, and a confidence score. These outputs together define a minimal edit to the original DiffPROTACs sample that repairs connectivity while preserving chemically reasonable geometry.

#### Input Encoding

The input is a complete molecule graph, where nodes are all the atoms in the PROTAC and the edges are constructed from the 3D coordinates by first building a k-nearest neighbors’ graph (https://www.geeksforgeeks.org/machine-learning/k-nearest-neighbours/) and filtering the neighbours with distance based bonding heuristics. Node features are a concatenated vector including a one hot atom type (with an additional KEEP class), binary masks marking the presence of linker and anchor atoms, and auxiliary scalars such as: distance to the closest anchor; soft connectivity or cutoff score; valence mismatch flag; and local graph degree. The 3D coordinates are stored separately as a per atom vector and are used to derive edge level geometric features: interatomic distances are expanded with RBF embeddings and combined with distance-based bonding signals, and these precomputed edge features are then fed into the attention layers so that message passing remains strongly geometry aware.

#### Inside GAT Layers

Inside each Pro-GAT GAT layer, node embeddings are updated by a multi-head, geometry-aware attention mechanism operating on the molecular graph [18, 21, 22].

Given node features 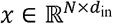 and edge features 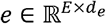, the layer first computes per-head query, key, value, and edge projections

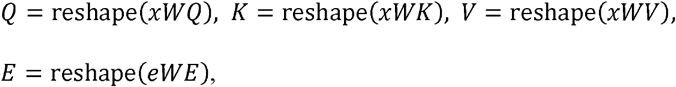

Where 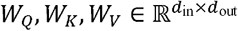 and 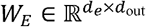, and the reshape splits the output into *H* heads of dimension 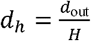.

For each directed edge *i* ← *j*, the attention score in head *h* is

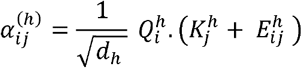

which combines the query of atom *i*, the key of neighbour *j*, and the learned edge embedding that encodes the RBF-expanded distance and unit bond vector [23, 24]. Normalization is done with a softmax over all incoming edges of a node

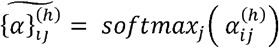

so that spatially and chemically compatible neighbours receive higher weight. The message from neighbour *j* to node *i* is then

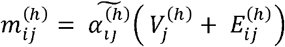

and the updated representation of node *i* is obtained by summing messages over all neighbours and heads,

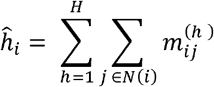

A final linear projection and residual-norm block produce the layer output

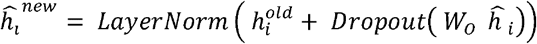

where 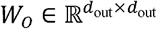.

Stacking several such layers allows each atom to iteratively accumulate information from an increasingly larger neighbourhood, so that its final embedding reflects local bonding geometry, long-range interactions across the linker and both anchors, and connectivity cues encoded in the edge features, before being passed to the type and coordinate MLP heads.

#### Node Embeddings before MLP

Each atom of the Protac has a final embedding vector that integrates its rich input features, such as element type, KEEP flag, linker/After several attention-based graph transformer [25] layers over the full anchor role, distance to anchors, soft connectivity score, valence mismatch, and local degree, together with multi-hop context from its neighbours and long-range interactions mediated by attention. In other words, this embedding reflects local chemistry (immediately bonding pattern and valence), global PROTAC topology (position within the linker relative to both anchors), and geometric cues injected through distance-derived edge features accumulated across layers [26].

#### MLP

The final node embeddings are fed to two parallel MLP heads: a type prediction head and a coordinate regression head [27].

The type head is a feed forward network with hidden size equal to the embedding dimension, followed by ReLU, dropout and finally a linear layer that outputs logits over the K atom types plus a dedicated KEEP class, 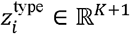 for each atom *i*.The coordinate head has the same structure but ends in a 3-dimensional linear layer with a tanh [28] activation, producing bounded coordinate offsets

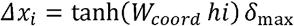

where δ_max_ = 0.06 Å, which are applied per atom to repair broken connectivity.

#### Output Processing

Type logits from the type prediction head are softmaxed to get per-atom probabilities over the 9 atom types (C, O, N, F, S, Cl, Br, I, P) plus a special ‘KEEP’ class (index 9), allowing the model to retain original atoms when minimal changes suffice.

Training utilizes class-balanced cross-entropy loss with upweights of 1.8× for heteroatoms (N, O, S, P), and 0.3× downweight for KEEP, which biases predictions toward chemically plausible edits such as C_→_N or C_→_O in linker regions while discouraging unnecessary changes.

During inference, a tuneable keep bias subtracts from KEEP logits and adds -1.5 to non-KEEP logits, dynamically controlling edit aggressiveness across multiple attempts.

Coordinate deltas are tanh scaled (max 0.6 Å) and filtered to only predictions with a norm greater than 0.05 Å, so only meaningful displacements are considered.

For edit candidates, calculate a composite priority (linker mask 2.0×, soft-cut score 1.5× measuring bottleneck position, and change probability 1 - keep prob), then sort descending and select the top-4 per iteration.

Type changes update atom logits to one-hot, while coordinate shifts apply at 0.5× magnitude; connectivity is verified post-edits via soft adjacency propagation, iterating up to 4 attempts or fallback heuristics.

#### Post Processing

Candidate PROTACs following Pro-GAT repair undergo a two-stage post-processing pipeline, combining RDKit sanitization with rule-based chemical filtering.

In Stage 1, RDKit (https://www.rdkit.org/) sanitization is applied and explicit valence errors are corrected using only local, geometry-preserving edits: reduction of excessive bond orders, removal of a minimal number of bonds and rebuilding of connectivity from 3D coordinates when needed, using covalent radius thresholds.

In Stage 2, RDKit derived descriptors are used to discard obviously problematic structures [29], including molecules with very few heavy atoms, reactive halides (Br/Cl/I), strongly charged alkoxides or non-neutral overall charge, and suspicious sulphur environments. Only molecules that pass both stages are retained for reporting validity, uniqueness, and related metrics.

An additional refinement step for retained molecules involves the selective application of MMFF-based energy minimization [30] on the linker atoms only, while keeping the ligand anchor geometries fixed. The targeted relaxation removes local strain introduced during the repair process without perturbing the global ligand geometry, so that downstream evaluations are carried out on physically reasonable conformations without confounding the actual repair process itself.

#### Hyperparameter configuration

The hyperparameters used in this work control the process of molecular graph generation, the structure of the graph attention network, and the training process. They include parameters associated with molecular graph generation and geometric representation, the depth and dimensionality of the model, attention mechanisms, and training parameters like the optimizer, learning rate, and loss functions. All these hyperparameters interact to ensure expressiveness, training stability, and chemical validity while facilitating molecular repair. The complete list of hyperparameters and their values is included in the supplementary Information [Table S1].

### 1.4. Evaluation Metrics

Our evaluation metrics are: validity, uniqueness, recovery, drug-likeness [31] and Synthetic Accessibility [32]. To evaluate model performance, the base diffusion model generated 10 samples per input ligand pair from the test set.

Disconnected samples underwent repair processing, yielding a final set of valid PROTAC conformations for aggregate metric computation.

#### Validity

Molecular validity determines if the generated PROTACs fall into chemically plausible space. This metric comprises: OpenBabel derived bond connectivity [21982300], RDKit valency rule compliance, no detached atoms given fixed input fragments and preservation of specified ligand components.

#### Uniqueness

Uniqueness is quantified as the proportion of non-redundant molecules among all valid generations. For each input *i*, canonical SMILES are computed in order to collapse molecules that are identical in 2D representation, which avoids the ambiguity of assessing equality in 3D space. The overall uniqueness score is then obtained as:

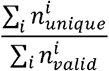

Where, 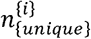 and 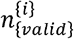 are the number of unique SMILES and the number of valid SMILES generated molecules for input *i*

#### Recovery Rate

The recovery rate evaluates the repair capability of Pro-GAT by measuring how many disconnected PROTACs produced by the diffusion model are successfully converted into connected structures after repair. It is computed as the fraction of disconnected diffusion outputs that are connected following Pro-GAT processing, relative to the total number of disconnected molecules before repair.

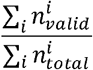

## 2. Results and Discussion

To contextualize the performance of Pro-GAT, we evaluate it as an explicit chemical validity stage applied to diffusion-generated PROTAC linker candidates for fixed POI–E3 ligand pairs from the held-out PROTAC-DB [33] test set used in DiffPROTACs. All reported metrics reflect aggregate output after enforcing chemical connectivity with Pro-GAT, rather than direct comparisons between standalone generative models.

We sample 10 candidate linkers for each anchor-ligand pair using a diffusion-based linker generator and compute chemical validity and uniqueness before and after enforcing chemical connectivity with Pro-GAT under identical conditions. This setup isolates the contribution of Pro-GAT in enforcing chemical realizability of diffusion-generated outputs without retraining or modifying the underlying generator.

We evaluate Pro-GAT across diffusion-based PROTAC generators, including DiffPROTACs and DiffLinker fine-tuned on PROTAC-DB [34] and presented as Table 1.

**Table 1.** Performance metrics of different methods on PROTAC test dataset.

| Method | Validity | Uniqueness | Recovery Rate |
| --- | --- | --- | --- |
| DiffPROTACs | 76.700% | 76.364% | 0% |
| DiffPROTACs + Pro-GAT | **83.918%** | **80.178%** | **31%** |
| DiffLinker-finetuned | 63.165% | 63.878% | 0% |
| DiffLinker-finetuned + Pro-GAT | 68.735% | 62.805% | 15.12% |

Table 1. summarizes performance of all the baselines on the shared PROTAC-DB test set in terms of validity, uniqueness and recovery rate. Bolded percentages indicate the best score of each metric.

Pro-GAT was trained on all ligand pairs from the PROTAC□DB training set but only on examples with linker lengths between 5 and 8 atoms, which were obtained by first generating connected linkers with DiffPROTACs and then constructing disconnected training inputs from these molecules. Despite this narrow linker-length regime during training, Pro-GAT maintains strong performance even for substantially larger linker lengths at test time, whereas the baseline models degrade rapidly. DiffLinker, in particular, performs poorly for long linkers even after fine-tuning on PROTAC □ DB, as illustrated by the sharp drop in its metrics for larger anchor distances shown in Figure 2.

**Figure 1.**
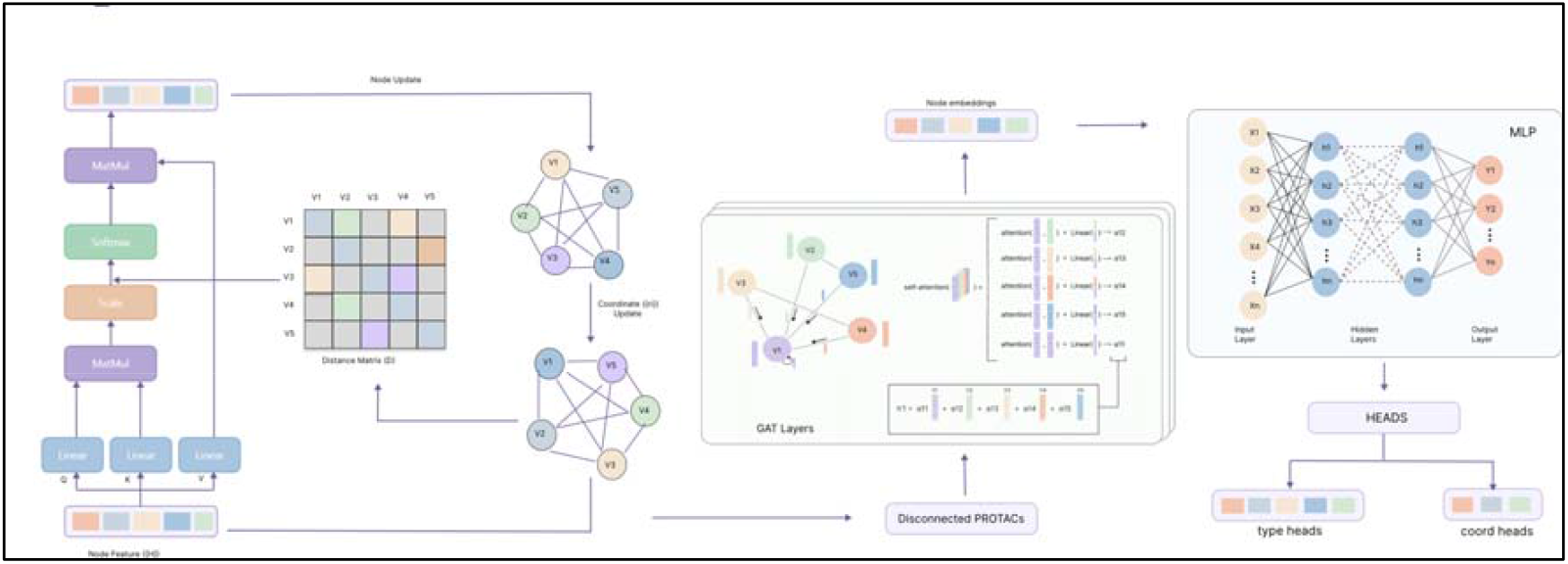
Overview of the Pro-GAT repair model pipeline with diffusion model’s architecture. Disconnected PROTACs are first converted into molecular graphs, processed through graph-attention layers, and decoded via MLP heads into atom-type edits and coordinate corrections to produce connected linkers.

**Figure 2.**
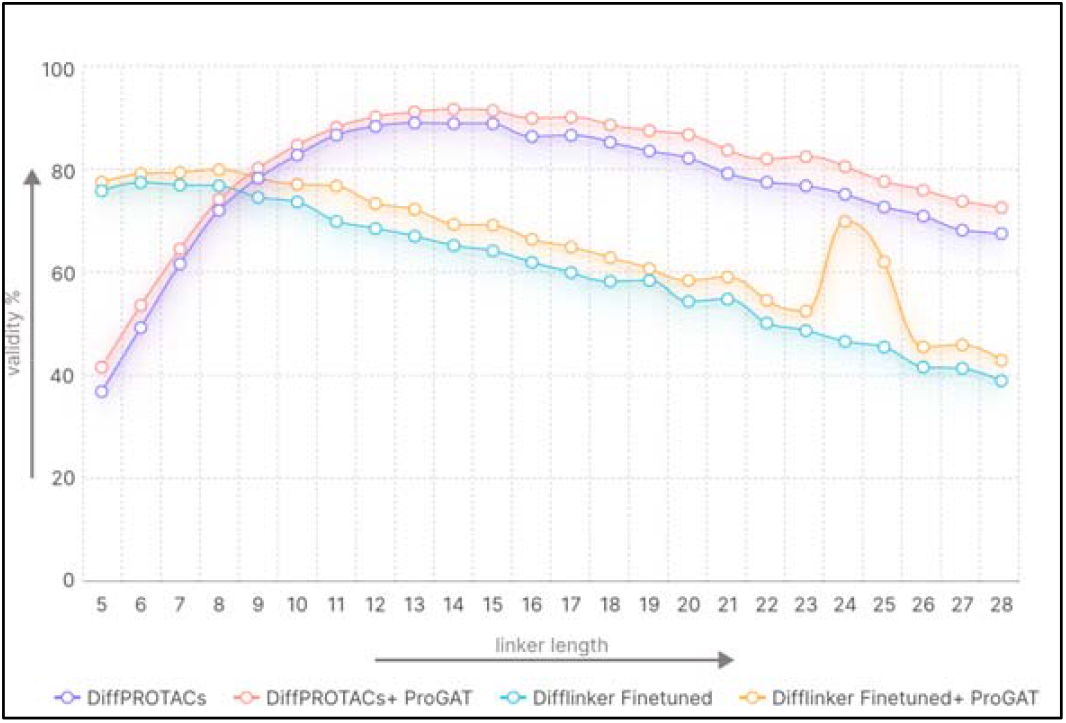
Effect of Pro-GAT on validity across linker lengths.

Figure 2. represents the validity (%) across linker lengths ranging from 5 to 28 atoms for DiffPROTACs and DiffLinker Finetuned with and without Pro-GAT. Pro-GAT improves the validity for both methods across the entire range. DiffPROTACs with Pro-GAT gives the highest overall validity. DiffLinker Finetuned also exhibits improved validity with Pro-GAT.

To evaluate whether Pro-GAT preserves geometric compatibility during the repair process, we analyze the magnitude and locality of the structural perturbations introduced during connectivity restoration. Pro-GAT explicitly constrains coordinate updates, bounding atomic displacements to at most 0.8 Å along each coordinate dimension (corresponding to an upper bound of approximately 1.4 Å in Euclidean space). These bounds prevent large-scale rearrangements of the diffusion-generated three-dimensional structure and ensure that all modifications remain strictly local to the disconnection region.

Within this more constrained regime, Pro-GAT reconnects covalently by locally adjusting coordinates and reassigning atom type without adding new atoms, extending linker length, or performing global conformational optimization. Any residual valence inconsistencies introduced in reconnection are eliminated deterministically through RDKit sanitization applied post-repair. The global spatial configuration output from the upstream diffusion model is thereby preserved across the repair stage.

### Structural quality of repaired linkers

Since Pro-GAT employs bounded local corrections without complete conformational optimization, bond lengths are the most indicative feature of any potential distortion introduced by the repair. To assess the robustness of the repaired linkers, we examine the bond length statistics for all linker bonds that appear at least 50 times in the 5,097 repaired PROTAC structures.

For each type of bond that qualifies, we examine the empirical distribution of bond distances in the repaired set. The full set of bond length statistics, by bond type, is provided in the <u>Supplementary Information</u>. The aim is to determine whether Pro-GAT maintains realistic geometry in the repaired structure, rather than optimizing ideal or force-field-adjusted bond lengths.

In general, the repaired linkers have a stable and chemically reasonable bond length distribution for all bond types, indicating that the local repairs by Pro-GAT do not introduce systematic geometric errors. Instances of pronounced deviation are rare and are predominantly associated with inputs that already exhibit globally unrealistic or highly strained geometries. In these situations, the geometric errors are not in the region of the disconnected linker bridge targeted by the repair process, and therefore fall beyond the scope of Pro-GAT’s bounded, locality-preserving corrections.

From these results, it appears that Pro-GAT preserves the integrity of the structure in the areas where it is intended to repair, without causing any undesired geometric distortions, even in the presence of difficult input geometries.

### Case Study: Geometry-Preserving Repair in a Known Ternary Complex

To assess if post-generation repair maintains the usability of the downstream structure, we perform a qualitative case study on a solved ternary complex (PDB ID: 7Z76) [35]. Starting from the crystallographic structure, the native PROTAC is isolated and broken down into its component ligands and linker. The ligand orientations are set to remain fixed as they are in the crystal structure to isolate the effect of linker regeneration.

With the help of DiffPROTACs, linker candidates of the same length as the original linker and fixed ligand geometry are produced. Among the ten samples generated, a chemically invalid structure is chosen and then corrected using Pro-GAT. The correction is done in a manner that only local modifications are allowed.

Following repair, the regenerated PROTAC is reinserted into the ternary complex after removal of the native molecule and is redocked [36] into the protein pair. Structural comparison indicates that the repaired PROTAC maintains overall molecular geometry and ligand poses (Fig. 3), with the regenerated linker spanning the original attachment points without introducing steric clashes or disrupting the protein–protein interface. Interaction analysis (Fig. 4) further shows that key ligand anchor interactions on both proteins are preserved, while differences are largely confined to linker-mediated contacts, consistent with the intrinsic flexibility of PROTAC linkers. Consistent with these observations, the repaired PROTAC achieves a docking score of −11.6, compared to −12.9 for the crystallographic reference.

**Figure 3.**
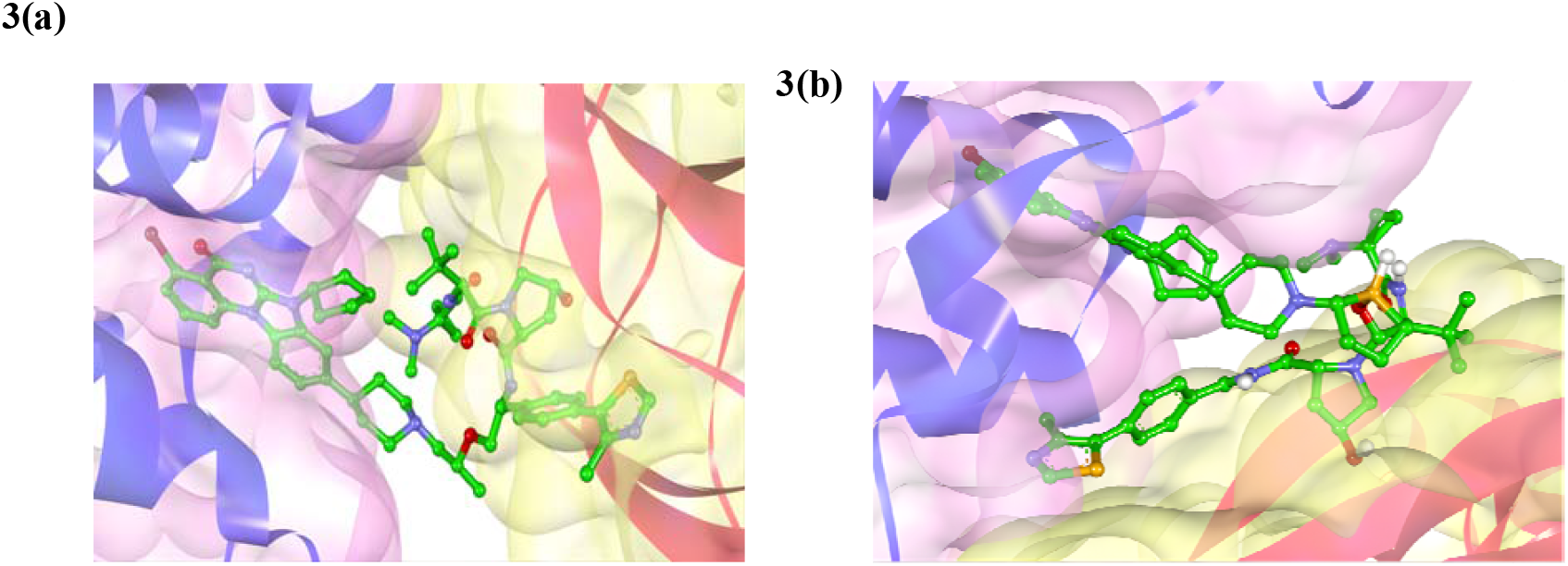
Comparison of the native and repaired PROTACs in the 7Z76 ternary complex by structural analysis. **(a)**Crystallographic PROTAC extracted from the 7Z76 ternary complex, shown in its native bound conformation with the E3 ligase and target protein. **(b)**Repaired PROTAC produced after linker regeneration and Pro-GAT-based repair, reinserted into the ternary complex after the removal of the native molecule.

**Figure 4.**
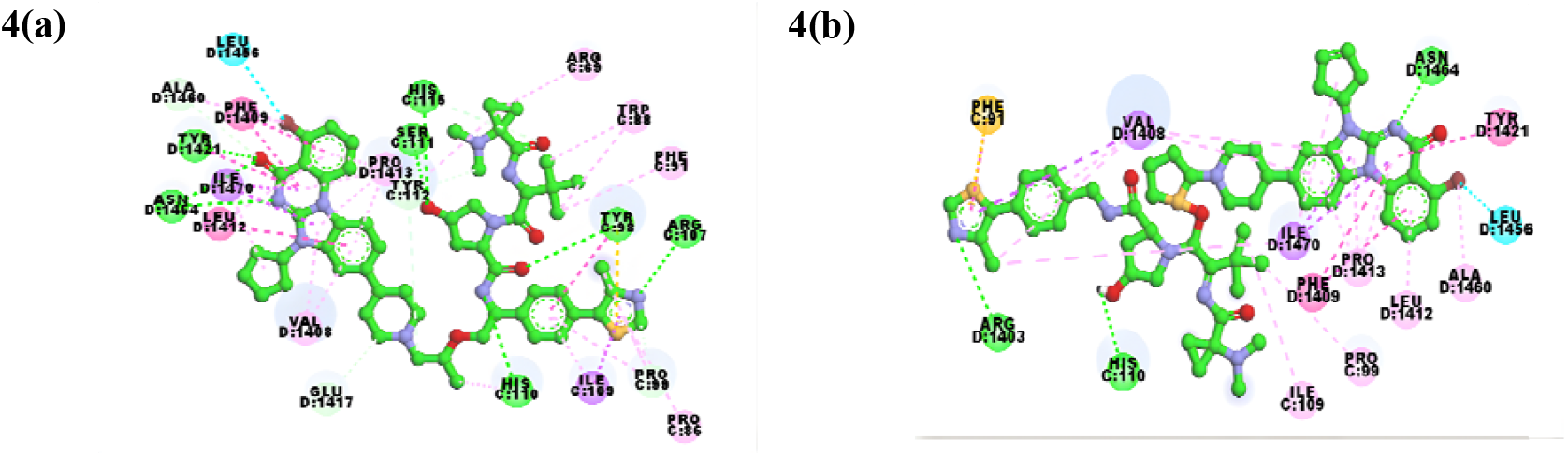
Comparison of protein–PROTAC interaction patterns in the 7Z76 ternary complex. **(a)**Protein–PROTAC interaction diagram for the crystallographic PROTAC. **(b)**Protein–PROTAC interaction diagram for the repaired PROTAC after linker regeneration and Pro-GAT–based repair.

## 3. Ablation Study

The generation of PROTACs was tested under three settings: diffusion baseline, diffusion with GAT repair module, and the full pipeline with GAT and post-processing. Across these ablations, the addition of GAT improved both validity and uniqueness, suggesting that modeling local graph structure in the generator is beneficial for enforcing chemical validity. The addition of the post-processing step resulted in the best-performing pipeline across all settings, achieving the highest validity and uniqueness of the generated chemicals. These findings suggest that the full pipeline outperforms the diffusion baseline in terms of aggregate output quality without sacrificing diversity.

Figure 5. illustrates the ablation of diffusion, diffusion with GAT, and diffusion with GAT+ post-processing on PROTAC validity, uniqueness, and recovery rate, which demonstrate the consistent improvement from Pro-GAT repair and filtering.

## 4. Conclusion

In present work, we propose Pro-GAT, a graph-based post-generation repair model that aims to enhance the number of useful outputs produced by diffusion-based PROTAC generators by repairing chemically disconnected outputs through local and geometry-preserving transformations. Rather than conflating geometric sampling with chemical correctness, Pro-GAT explicitly separates exploration from realizability, addressing a fundamental limitation of continuous generative models for molecular design. This is more concerned with chemical disconnection, structural correctness, and usability

**Figure 5.**
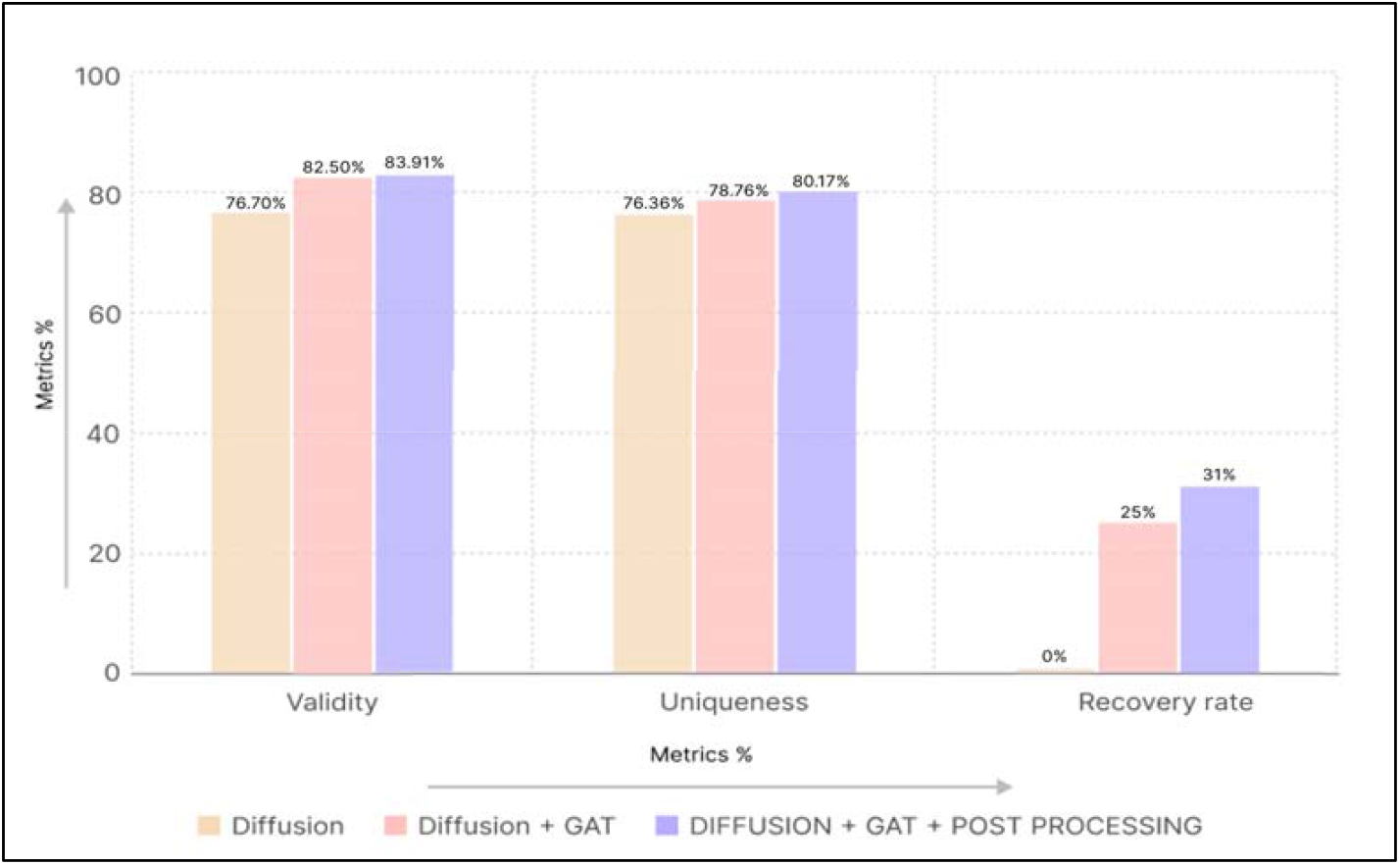
Ablation study of Pro-GAT-based post-processing on PROTAC generation quality metrics

Our results indicate that Pro-GAT is successful in repairing a large number of otherwise abandoned PROTACs, leading to a substantial increase in overall validity with realistic sampling budgets, while maintaining high uniqueness and small geometric perturbation. Most importantly, the network intentionally decides to abstain from repairing those that are, in principle, impossible to repair, such as those arising from a short linker or a large anchor-point distance, thereby ensuring that physical reality is preserved and no connectivity is hallucinated. This is a reflection of a strict division of responsibilities between generation and repair, enabling diffusion models to focus on exploration while delegating chemical correctness to a learned post-processing stage.

From a practical perspective, Pro-GAT addresses a critical but underexplored bottleneck in structure-based PROTAC design: the large fraction of invalid structures produced by generative models that are silently filtered out in current pipelines. By recovering thousands of chemically valid candidates without retraining or modifying the underlying generator, Pro-GAT improves sample efficiency, reduces computational waste, and enables downstream tasks such as docking, molecular dynamics, and ternary complex analysis to be applied to a broader and more representative set of generated molecules.

Although Pro-GAT is evaluated with DiffPROTACs and DiffLinker, the proposed repair approach is model-agnostic and can, in principle, be integrated with any 3D molecular generator that produces graphs with coordinates. Future work could explore the extension of the repair problem to incorporate task-specific constraints, such as binding site topology or physicochemical property preservation, as well as the integration of Pro-GAT with generation/repair loops. More broadly, this work highlights the importance of explicitly modeling post-generation validity as a first-class problem in molecular machine learning, and provides a practical step toward more reliable and usable generative design pipelines for PROTACs and other complex molecular modalities.

## DECLARATIONS

### Ethics approval and consent to participate

Not applicable

### Consent for publication

Not applicable

### Supplementary Material

Supplementary information and tables are provided as Supplementary information File.

### Data availability

All data supporting the findings of this study are available in the Supplementary Materials as comma-separated (.csv) files.

### Author Contributions

**S.V**.: Investigation; validation; methodology; visualization; writing—original draft; formal analysis. **L.P.B**.: Data curation; formal analysis; methodology; validation; visualization. **S.G**.: Conceptualization; methodology; validation; visualization; formal analysis; data curation; software; supervision; formal analysis; writing—review and editing. **V.K**.: Conceptualization; methodology; writing—review and editing; supervision; formal analysis.

## Acknowledgements

We authors, Saahithi Vemuri, Laxmi Priya Bijigiri, Sanjana Gogte, and Vani Kondaparthi, express our sincere gratitude to the Drugparadigm Research Lab for providing the necessary facilities and infrastructure that enabled the successful completion of this work.

## REFERENCES

1. Guedeney N, Cornu M, Schwalen F, Kieffer C, Voisin-Chiret AS. PROTAC technology: A new drug design for chemical biology with many challenges in drug discovery. Drug Discov Today. [Research Support, Non-U.S. Gov’t Review]. 2023;28(1):103395. DoI:10.1016/j.drudis.2022.103395

2. Cecchini C, Tardy S, Ceserani V, Theurillat JP, Scapozza L. Exploring the Ubiquitin-Proteasome System (UPS) through PROTAC Technology. Chimia (Aarau). 2020;74(4):274–7. DoI:10.2533/chimia.2020.274

3. Chen Z, Gu C, Tan S, Wang X, Li Y, He M, et al. Interpretable PROTAC Degradation Prediction With Structure-Informed Deep Ternary Attention Framework. Adv Sci (Weinh). 2025;12(47):e08138. DoI:10.1002/advs.202508138

4. M S, Joga R, Gandhi K, Yerram S, Raghuvanshi RS, Srivastava S. Exploring the clinical trials, regulatory insights, and challenges of PROTACs in oncology. Semin Oncol. [Review]. 2025;52(2):152339. DoI:10.1016/j.seminoncol.2025.152339

5. Berkley K, Zalejski J, Sharma N, Sharma A. Journey of PROTAC: From Bench to Clinical Trial and Beyond. Biochemistry. [Review]. 2025;64(3):563–80. DoI:10.1021/acs.biochem.4c00577

6. Ibrahim S, Umer Khan M, Khurram I, Rehman R, Rauf A, Ahmad Z, et al. Navigating PROTACs in Cancer Therapy: Advancements, Challenges, and Future Horizons. Food Sci Nutr. [Review]. 2025;13(2):e70011. DoI:10.1002/fsn3.70011

7. Li X, Song Y. Proteolysis-targeting chimera (PROTAC) for targeted protein degradation and cancer therapy. J Hematol Oncol. [Research Support, Non-U.S. Gov’t Research Support, U.S. Gov’t, Non-P.H.S. Review]. 2020;13(1):50. DoI:10.1186/s13045-020-00885-3

8. Dong Y, Ma T, Xu T, Feng Z, Li Y, Song L, et al. Characteristic roadmap of linker governs the rational design of PROTACs. Acta Pharm Sin B. [Review]. 2024;14(10):4266–95. DoI:10.1016/j.apsb.2024.04.007

9. Mostofian B, Martin HJ, Razavi A, Patel S, Allen B, Sherman W, et al. Targeted Protein Degradation: Advances, Challenges, and Prospects for Computational Methods. J Chem Inf Model. [Review]. 2023;63(17):5408–32. DoI:10.1021/acs.jcim.3c00603

10. Yang Q, Zhao J, Chen D, Wang Y. E3 ubiquitin ligases: styles, structures and functions. Mol Biomed. [Review]. 2021;2(1):23. DoI:10.1186/s43556-021-00043-2

11. Samarasinghe KTG, Crews CM. Targeted protein degradation: A promise for undruggable proteins. Cell Chem Biol. [Research Support, N.I.H., Extramural Research Support, Non-U.S. Gov’t Review]. 2021;28(7):934–51. DoI:10.1016/j.chembiol.2021.04.011

12. Li F, Hu Q, Zhou Y, Yang H, Bai F. DiffPROTACs is a deep learning-based generator for proteolysis targeting chimeras. Brief Bioinform. 2024;25(5). DoI:10.1093/bib/bbae358

13. Ho J, Jain A, Abbeel P. Denoising Diffusion Probabilistic Models. ArXiv. 2020;abs/2006.11239

14. Song Y, Sohl-Dickstein JN, Kingma DP, Kumar A, Ermon S, Poole B. Score-Based Generative Modeling through Stochastic Differential Equations. ArXiv. 2020;abs/2011.13456

15. Brown N, Fiscato M, Segler MHS, Vaucher AC. GuacaMol: Benchmarking Models for de Novo Molecular Design. J Chem Inf Model. 2019;59(3):1096–108. DoI:10.1021/acs.jcim.8b00839

16. Ge J, Li S, Weng G, Wang H, Fang M, Sun H, et al. PROTAC-DB 3.0: an updated database of PROTACs with extended pharmacokinetic parameters. Nucleic Acids Res. 2025;53(D1):D1510–D5. DoI:10.1093/nar/gkae768

17. Troup RI, Fallan C, Baud MGJ. Current strategies for the design of PROTAC linkers: a critical review. Explor Target Antitumor Ther. [Review]. 2020;1(5):273–312. DoI:10.37349/etat.2020.00018

18. Velickovic P, Cucurull G, Casanova A, Romero A, Liò P, Bengio Y. Graph Attention Networks. ArXiv. 2017;abs/1710.10903

19. Rumelhart DE, Hinton GE, Williams RJ. Learning representations by back-propagating errors. Nature. 1986;323(6088):533–6. DoI:10.1038/323533a0

20. LeCun YA, Bottou L, Orr GB, Müller K-R. Efficient BackProp. In: Montavon G, Orr GB, Müller K-R, editors. Neural Networks: Tricks of the Trade: Second Edition. Berlin, Heidelberg: Springer Berlin Heidelberg; 2012. p. 9–48.

21. Xu L, Pan S, Xia L, Li Z. Molecular Property Prediction by Combining LSTM and GAT. Biomolecules. [Research Support, Non-U.S. Gov’t]. 2023;13(3). DoI:10.3390/biom13030503

22. Vaswani A, Shazeer N, Parmar N, Uszkoreit J, Jones L, Gomez AN, et al., editors. Attention is All you Need. Neural Information Processing Systems; 2017.

23. Gilmer J, Schoenholz SS, Riley PF, Vinyals O, Dahl GE, editors. Neural Message Passing for Quantum Chemistry. International Conference on Machine Learning; 2017.

24. Battaglia PW, Hamrick JB, Bapst V, Sanchez-Gonzalez A, Zambaldi VF, Malinowski M, et al. Relational inductive biases, deep learning, and graph networks. ArXiv. 2018;abs/1806.01261

25. Shehzad A, Xia F, Abid S, Peng C, Yu S, Zhang D, et al. Graph Transformers: A Survey. IEEE transactions on neural networks and learning systems. 2024;PP

26. Satorras VG, Hoogeboom E, Welling M, editors. E(n) Equivariant Graph Neural Networks. International Conference on Machine Learning; 2021.

27. Liu R, Li Y, Tao L, Liang D, Zheng HT. Are we ready for a new paradigm shift? A survey on visual deep MLP. Patterns (N Y). [Review]. 2022;3(7):100520. DoI:10.1016/j.patter.2022.100520

28. De Ryck T, Lanthaler S, Mishra S. On the approximation of functions by tanh neural networks. Neural Netw. 2021;143:732–50. DoI:10.1016/j.neunet.2021.08.015

29. Li Q, Liu C, Shashikumar SP, Nemati S, Shen Z, Clifford GD. Ventricular ectopic beat detection using a wavelet transform and a convolutional neural network. Physiol Meas. [Research Support, N.I.H., Extramural Research Support, Non-U.S. Gov’t Research Support, U.S. Gov’t, Non-P.H.S.]. 2019;40(5):055002. DoI:10.1088/1361-6579/ab17f0

30. Halgren TA. MMFF VI. MMFF94s option for energy minimization studies. J Comput Chem. 1999;20(7):720–9. DoI:10.1002/(SICI)1096-987X(199905)20:7<720::AID-JCC7>3.0.CO;2-X

31. Bickerton GR, Paolini GV, Besnard J, Muresan S, Hopkins AL. Quantifying the chemical beauty of drugs. Nat Chem. [Research Support, Non-U.S. Gov’t]. 2012;4(2):90–8. DoI:10.1038/nchem.1243

32. Ertl P, Schuffenhauer A. Estimation of synthetic accessibility score of drug-like molecules based on molecular complexity and fragment contributions. J Cheminform. 2009;1(1):8. DoI:10.1186/1758-2946-1-8

33. Weng G, Cai X, Cao D, Du H, Shen C, Deng Y, et al. PROTAC-DB 2.0: an updated database of PROTACs. Nucleic Acids Res. [Research Support, Non-U.S. Gov’t]. 2023;51(D1):D1367–D72. DoI:10.1093/nar/gkac946

34. Igashov I, Stärk H, Vignac C, Schneuing A, Satorras VG, Frossard P, et al. Equivariant 3D-conditional diffusion model for molecular linker design. Nature Machine Intelligence. 2024;6(4):417–27. DoI:10.1038/s42256-024-00815-9

35. Berman HM, Westbrook J, Feng Z, Gilliland G, Bhat TN, Weissig H, et al. The Protein Data Bank. Nucleic Acids Res. [Research Support, U.S. Gov’t, Non-P.H.S. Research Support, U.S. Gov’t, P.H.S.]. 2000;28(1):235–42. DoI:10.1093/nar/28.1.235

36. Eberhardt J, Santos-Martins D, Tillack AF, Forli S. AutoDock Vina 1.2.0: New Docking Methods, Expanded Force Field, and Python Bindings. J Chem Inf Model. [Research Support, N.I.H., Extramural]. 2021;61(8):3891–8. DoI:10.1021/acs.jcim.1c00203

