## Supplementary Information for "Pro-GAT: Reconnecting Fragmented PROTACs Using Graph Attention Transformer"

The file containing Supplementary Information is divided into two parts:

(1) Hyperparameter tuning of Pro-GAT, including data preprocessing and feature definition, model architecture selection, training parameters, and loss function definition.

(2) Bond Length Statistics for Repaired PROTAC Linkers, which provides the quantitative bond length distribution for the repaired linkers and a description of typical failure examples to illustrate the extent of the local repair method.

In addition to examples of geometric infeasibility on a global scale, where the distance between ligand binding points is greater than the existing geometry of the linker backbone can span, Pro-GAT may also fail when the restoration of chemical connectivity necessitates non-local topological transformations that are beyond the capabilities of the local repair algorithms used by the model. Examples of this include cases that necessitate large coordinate translations or atom insertions, which are beyond the capabilities of Pro-GAT. Finally, the repair problem is defined with a soft, differentiable dependence on connectivity functions that approximate reachability during optimization. While this is helpful for guiding the optimization process, it can sometimes result in cases where soft connectivity is achieved but a discrete, chemically valid bond graph cannot be constructed according to strict valence constraints and RDKit sanitization.

### 1. Hyperparameter configuration

Atoms are parameterized as nodes with identity logits and additional connectivity features, while molecular structure is represented as a k-nearest-neighbour graph in three-dimensional space. Edge features capture interatomic distances using radial basis functions combined with relative direction information, allowing the model to reason about molecular geometry. Binary anchor and linker masks are used as guidance to distinguish fixed atoms from atoms that can be modified.

The model uses a graph attention network with multiple layers and heads to spread information throughout the molecular graph. Attention is used to weight neighbouring atoms based on relevance, and a fixed hidden dimension is used for expressive but stable representations. The output heads predict atom type edits (with a KEEP option for unchanged atoms) and bounded coordinate changes, making it easier to make conservative and targeted edits.

The training is done at the molecular level, using individual molecules, with the AdamW optimizer and gradient clipping. The loss function combines atom type classification, coordinate regression, connectivity enforcement, valence enforcement, and sparsity. These loss functions combined help the model predict the connectivity of molecules while being chemically valid and avoiding unnecessary changes.

**Table S1. Summary of Model Hyperparameters**

| Category | Hyperparameter | Value |
|---|---|---|
| Data & Features | Atom Types | 9(C, O, N, F, S, Cl, Br, I, P) |
| | Node Feature Dimension | Atom logits + 6 auxiliary features |
| | Anchor Mask | Binary |
| | Linker Mask | Binary |
| | k-NN graph (k) | 10 |
| | RBF Kernels | 16 |
| | RBF Distance Cutoff | 8.0 Å |
| | Edge Feature Dimension | 19 (16 RBF + 3 direction) |
| Model Architecture | Model Type | Graph Attention Network |
| | Hidden Dimension | 256 |
| | Number of Graph Layers | 6 |
| | Attention Heads | 4 |
| | Node Embedding Dimension | 256 |
| | Edge Embedding Dimension | 128 |
| | Output classes | 10 (9 atom types + KEEP) |
| Training | Optimizer | AdamW |
| | Learning Rate | 1e-3 |
| | Weight Decay | 1e-4 |
| | Batch Size | 1 |
| | Epochs | 200 |
| | Gradient Clip Norm | 1.0 |
| Loss Function | Atom Type Loss | Cross-Entopy |
| | Coordinate loss | Smooth L1 |
| | Connectivity Loss | Soft reachability |
| | Valence Loss | Degree mismatch |
| | Sparsity Loss | KEEP-biased edit minimization |

Table S1 lists the key model architecture and training parameters, along with their corresponding values, used in the development of the proposed molecular repair model.

### 2. Bond Length Statistics for Repaired PROTAC Linkers

To determine whether Pro-GAT maintains chemically plausible local geometry in linker repair, we carried out a systematic bond length analysis of repaired PROTACs and compared the distributions to reference values. For each molecule, three-dimensional bond lengths were calculated directly from RDKit conformers, considering only bonds where both endpoints were entirely contained within the linker region targeted by the repair method. Bonds were categorized by atom pair and bond type (single, double, triple, or aromatic), and summary statistics including count, mean and median bond length, standard deviation, and extrema were calculated for all linker lengths and molecules. Reference bond lengths were assigned according to experimentally determined values, and asymmetric tolerance intervals were employed to detect bonds that were compressed or extended, accounting for the relative ease of bond extension versus compression. In addition to measures of central tendency and dispersion, deviation statistics reporting bonds lying outside conservative tolerance intervals and extreme values were also recorded.

The complete set of bond length statistics is given in two CSV files, one for DiffPROTACs structures(DiffPROTACs.csv) and one for Pro-GAT structures(Pro-GAT.csv). Over the set of bond types present with sufficient frequency, the repaired linkers show stable distributions of bond lengths around the reference values, suggesting that the bounded, local repair process of Pro-GAT preserves chemically plausible linker geometry without requiring ideal or force-field-optimized bond lengths.
